# Systematic tissue oxygen variation shows the modulation of murine skin radiation toxicity at ultra-high dose rates

**DOI:** 10.1101/2025.10.06.680759

**Authors:** David I. Hunter, Jacob P. Sunnerberg, Armin D. Tavakkoli, Austin M. Sloop, Beverly Petusseau, Jiang Gui, Xu Cao, Rongxiao Zhang, Simin Belali, Harold M. Swartz, Lesley A. Jarvis, P. Jack Hoopes, David J. Gladstone, Brian W. Pogue

## Abstract

**Objective:** This study evaluated the hypothesis that baseline tissue oxygen (pO_2_) would modulate FLASH toxicity sparing in murine skin, using a wide range of pO_2_ values, with ultra-high dose rate (UHDR) versus conventional dose rate (CDR) irradiation.

**Approach:** Murine leg tissue pO_2_ was systematically varied and measured during irradiation from a FLASH Mobetron 9 MeV linac at 25 Gy, comparing UHDR (≈240 Gy/s) to CDR (≈0.16 Gy/s), for radiation induced skin toxicity outcomes. Baseline tissue pO_2_ was systematically modulated in 5 different treatment cohorts, using different ranges of inhaled gas (room air, 100% oxygen, or carbogen) and through varying limb vascular compression (partial or full). Radiolytic oxygen consumption, g_O2_ (mmHg/Gy), was quantified *in vivo*, and induced macroscopic skin toxicity was scored daily post treatment.

**Main Results:** FLASH skin sparing was observed at a fixed dose of 25 Gy in groups with partial leg clamping (pO_2_≈7±4mmHg), inhaled air (pO_2_≈12±6mmHg) and 100% oxygen (pO_2_≈16±4mmHg), while reduction in ulceration progression was significant only in the air inhalation group. No FLASH effect was observed under anoxic conditions, via complete blood flow occlusion (pO_2_≈0±1mmHg), or when modulated by inhaled carbogen (pO_2_≈21±7mmHg). *In vivo* measurements of radiolytic oxygen consumption, g_O2_, correlated to initial pO_2_ under UHDR conditions (pO_2_≈4-16mmHg), with ulceration predominantly occurring at pO_2_ values above 16mmHg. Inspired carbogen induced the highest pO_2_ at which point there was no FLASH sparing, for any dose groups between 25 to 15 Gy, despite having large changes in damage with dose. At the specific dose level of 25 Gy studied, the toxicity scores under anoxia for both UHDR and CDR were low (toxicity scores < 1) with no differences observed.

**Significance:** These findings point to the fact that moderate and low tissue pO_2_ is associated with diminished oxygen-mediated damage at UHDR but not CDR, seen with inspired room air or 100% oxygen. Anoxic and hyperoxic murine skin are associated with minimal and maximal radiation damage respectively, but also exhibit no apparent FLASH toxicity sparing effect, with further investigation warranted into if the FLASH toxicity sparing effect persists at higher doses under anoxia.

## Introduction

Oxygen plays a fundamental role in radiation therapy as demonstrated in cell survival and *in vivo* studies where the absence of oxygen (anoxia) can dramatically reduce radiation killing efficacy by a factor of ≈2.7, known as the oxygen enhancement ratio (OER) (Hall and Amato, 2018). Oxygen is one of the most reactive molecules in biology, and its presence amplifies radiation effectiveness through complex reactive oxygen species (ROS) that react with nearly all biological molecules, including DNA, in a process that “fixes” radiation induced damage (Wardman, 2007). Studies have shown that oxygen acts as a modulator of the observed FLASH effect (Kirby-Smith and Dolphin, 1958; Dewey and Boag, 1959; Hornsey and Bewley, 1971; Petersson *et al*., 2020; Tavakkoli *et al*., 2024), a phenomenon characterized by reduced normal tissue toxicity under ultra-high dose rate (UHDR) radiation therapy, where dose rates surpass 40-100 Gy/s, while maintaining a tumor control achieved at conventional dose rates (CDR, ∼0.17 Gy/s) (Favaudon *et al*., 2014; Montay-Gruel *et al*., 2017). One of many key distinctions between tumors and healthy tissues lies in their baseline oxygenation, with solid tumors typically being hypoxic (0-10 mmHg) as compared to normal tissues (10-40 mmHg) (Vaupel, Flood and Swartz, 2021). Solid tumors also have a heterogeneous distribution of oxygen tensions throughout their volume, further differentiating them from normal tissues (Hall and Amato, 2018). This difference may be particularly relevant in the context of radiotherapy given the role that oxygen tension plays in promoting the oxidative fixation of DNA damage (Grimes and Partridge, 2015; Hall and Amato, 2018). Perhaps the most compelling aspect of this is that true pO_2_ values are rarely known for any tissue being irradiated, and yet the OER effect can dramatically increase or decrease the effective toxicity to the underlying tissue, from any given dose delivery, altering the therapeutic ratio of any particular treatment or a course of treatment. In this study, we examined how prospectively measuring and modulating the tissue pO_2_ might affect the observation of the FLASH effect.

One of the more interesting features of UHDR delivery is that the dose and radiolytic cascade of ROS induced DNA damage timelines are sufficiently faster than the oxygen supply from the capillaries, so that it is possible to directly measure the oxygen consumed *in vivo* during ROS production (Cao *et al*., 2021; El Khatib *et al*., 2024; Grilj *et al*., 2024; Sunnerberg *et al*., 2025). A series of studies have shown how to quantify radiolytic oxygen consumption under UHDR conditions, and in solutions the oxygen consumption per unit dose (g_O2_) at UHDR is lower than that at conventional dose rates (CDR) (Cao *et al*., 2021; Jansen *et al*., 2021; El Khatib *et al*., 2022; Van Slyke *et al*., 2022; Sunnerberg *et al*., 2023). This suggests that UHDR may induce less ROS than CDR at isodose levels and thereby potentially result in less normal tissue toxicity. However, these observations with optical probes have only been made in *in vitro* solutions, and CDR measurements of this effect *in vivo* are not readily possible.

*In vivo* studies have shown stable and repeatable murine tissue oxygen consumption rates (El Khatib *et al*., 2024; Grilj *et al*., 2024; Sunnerberg *et al*., 2025) that reach a plateau of ≈ 0.2 - 0.3 mmHg/Gy above a 20 mmHg baseline tissue oxygenation (Sunnerberg *et al*., 2025). This finding matches previous *in vitro* studies (El Khatib *et al*., 2022; Ha *et al*., 2022; Jansen *et al*., 2022; Van Slyke *et al*., 2022; Sunnerberg *et al*., 2023) showing that the oxygen consumption g-value (g_O2_) is linear with oxygen tension below 20 mmHg, with a 50% level at approximately 8 mmHg, and a constant g_O2_ at oxygen levels above 20 mmHg. Even more important than this is the observation that the apparent g_O2_ values appear lower in tumor tissue than normal tissues (Cao *et al*., 2021), suggesting that there may be a reduction in ROS mediated damage. While these studies establish a clear relationship between tissue oxygenation and radiation-induced oxygen consumption, the correlation between these radiation-chemical patterns and differential tissue response under UHDR conditions remains unexplored.

This study was developed to explore the relationships between the observation of a skin toxicity effect with UHDR versus CDR, with the factors of initial tissue oxygen, delivered radiation dose and g_O2_ consumption yield. While recent studies have extensively examined the normal tissue complication probability (NTCP) response with variation in dose (Hansen *et al*., 2026; Sesink *et al*., 2026), including extensive summations of the current literature by Colizzi et al. (Colizzi *et al*., 2026), this has been at fixed values of oxygenation conditions. The goal of this current work was to systematically vary tissue oxygen to determine how tissue radiation toxicity varies with tissue pO_2_ at a fixed dose level, and if there was a difference between pO_2_ and g_O2_ in categorizing this effect. While it is common to vary dose as a modulator of FLASH, it is considerably less studied how systematic pO_2_ or g_O2_ variation affects the toxicity sparing FLASH effect.

## Methods

### Animals

This vertebrate animal study protocol was approved by the institutional IACUC committee, and all procedures carried out complied. One-hundred and fifty male Albino B6 mice were purchased (Jackson Laboratories, Cambridge, MA) at 8-10 weeks of age and housed in the institution vivarium. The population was divided into two groups, for either UHDR or conventional dose rate (CDR), with each group further split into five oxygenation cohorts. Each cohort consisted of 10-11 mice. There is a well-known sex differential in radiation response (National Research Council (US) Committee on the Biological Effects of Ionizing Radiation (BEIR V), 1990; Hall and Amato, 2018), which has been documented in the FLASH effect and so for simplicity only male mice were studied here (Tavakkoli *et al*., 2024).

Fur on the left leg, flank, and surrounding area of each mouse was shaved 3 days prior to irradiation. One hour before irradiation, all mice underwent anesthesia using isoflurane at 3% for induction (delivered at 500 ml/min for 3 minutes) and maintained at 1.5% (delivered at 100 ml/min) throughout procedures. The oxygen probe Oxyphor PdG4 (Oxygen Enterprises, Philadelphia, PA), at 200µM in 0.05 mL in phosphate buffer solution was systemically administered into the retro-orbital space, with the oxygen measurement procedure described further below. Mice were allowed to recover to permit the diffusion of Oxyphor PdG4 into all tissues as in prior studies (Sunnerberg *et al*., 2026). The mice were then re-anesthetized for radiation delivery, with identical induction and maintenance conditions, under three different gas conditions: (i) room air (21% oxygen), (ii) 100% oxygen, or (iii) carbogen (95% oxygen, 5% CO_2_), and three different levels of vascular clamping: (i) no clamping, (ii) partial clamping and (iii) complete clamping. The anesthesia/gas inhalation timing followed the protocol reported by Tavakkoli et al. a heating pad was placed underneath each mouse to maintain consistent body temperature (Tavakkoli *et al*., 2024).

### Radiation Delivery & Dosimetry

Irradiations were performed using a UHDR-capable 9 MeV Mobetron linear accelerator (IntraOp Inc., Sunnyvale, CA) designed for research use. The electron radiation field was shaped by a custom-designed collimator creating a 1.6 cm diameter circular treatment area, positioned on each mouse’s left hind leg (thigh and lower leg area). To ensure accurate dosimetry and simultaneous oxygen measurements, the experimental setup incorporated a 3-D printed PETG cone with integrated holders for both an EDGE™ Detector (Sun Nuclear Corp, Melbourne, FL) and oxygen measurements via a fiber-coupled phosphorimeter. This arrangement positioned the measurement devices upstream of the mouse while maintaining field alignment and preventing beam perturbation. To replicate conditions known to induce a FLASH sparing effect, all mice received a single-fraction dose of 25 Gy, with carbogen breathing mice receiving 25, 20 and 15 Gy, to the left leg using a 1.6 cm circular field, under either CDR or UHDR delivery modes (Duval *et al*., 2023).

Dosimetric accuracy was ensured through calibration of an EDGE™ Detector against a flashDiamond detector (PTW, Freiburg, Germany), specifically designed for UHDR applications. The calibration was performed under identical geometric and delivery conditions as the animal irradiations, establishing a conversion factor between diode readings and delivered electron fluence. Both detectors have been validated for UHDR dosimetry in previous studies by Rahman et al. and Tessonnier et al. (Rahman *et al*., 2023; Tessonnier *et al*., 2024). Temporal pulse information was measured and verified using the Beam Current Transformers (BCTs) installed in the head of the Mobetron, with dosimetric measurements included in Table 1.

**Table 1:**
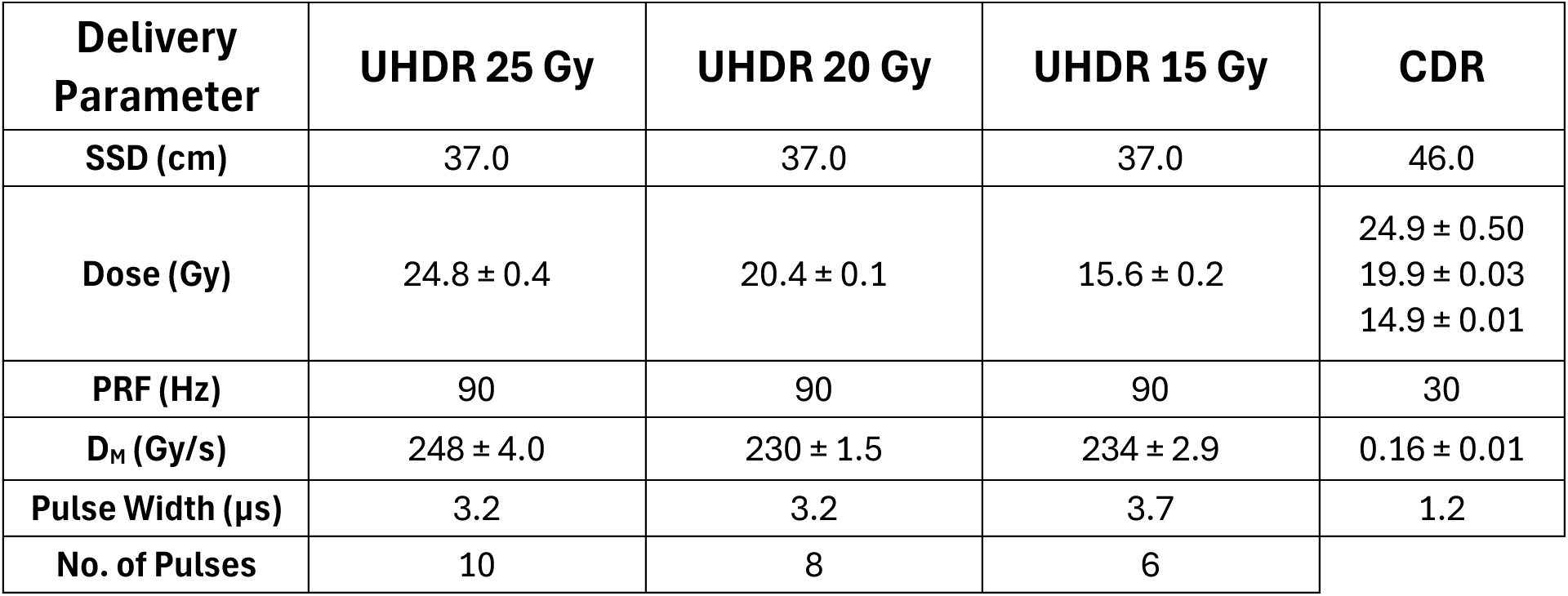
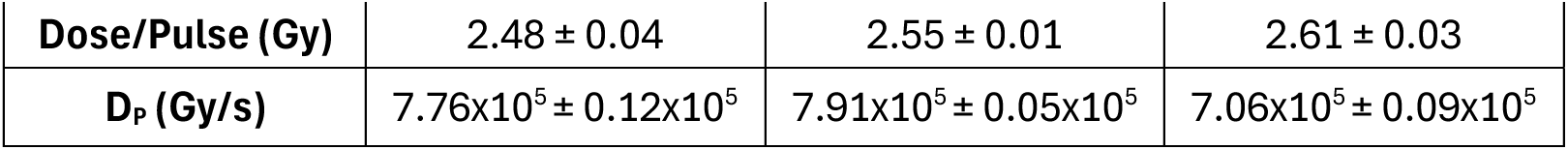
Beam delivery parameters for UHDR and CDR irradiations. Key characteristics include source-to-surface distance (SSD), pulse repetition frequency (PRF), pulse width, dose per pulse, instantaneous dose rate (D_P_), mean dose rate (D_M_), and total dose delivered, with values here reported as delivered with standard deviation values.

| **Delivery Parameter** | **UHDR 25 Gy** | **UHDR 20 Gy** | **UHDR 15 Gy** | **CDR** |
| --- | --- | --- | --- | --- |
| **SSD (cm)** | 37.0 | 37.0 | 37.0 | 46.0 |
| **Dose (Gy)** | 24.8 ± 0.4 | 20.4 ± 0.1 | 15.6 ± 0.2 | 24.9 ± 0.50<br>19.9 ± 0.03<br>14.9 ± 0.01 |
| **PRF (Hz)** | 90 | 90 | 90 | 30 |
| **D<sub>M</sub> (Gy/s)** | 248 ± 4.0 | 230 ± 1.5 | 234 ± 2.9 | 0.16 ± 0.01 |
| **Pulse Width (µs)** | 3.2 | 3.2 | 3.7 | 1.2 |
| **No. of Pulses** | 10 | 8 | 6 |  |

**Table 1:** Beam delivery parameters for UHDR and CDR irradiations.
|  |  |  |  |
| --- | --- | --- | --- |
| **Dose/Pulse (Gy)** | 2.48 ± 0.04 | 2.55 ± 0.01 | 2.61 ± 0.03 |
| **D<sub>P</sub> (Gy/s)** | 7.76x10<sup>5</sup> ± 0.12x10<sup>5</sup> | 7.91x10<sup>5</sup> ± 0.05x10<sup>5</sup> | 7.06x10<sup>5</sup> ± 0.09x10<sup>5</sup> |

### Radiation Induced Skin Toxicity

Following radiation delivery, animals were monitored daily to visually score the progression of radiation induced skin toxicity, via fixed geometry white light imaging. The primary endpoint of the study was the magnitude of the skin toxicity score at the irradiation site, with a secondary measure of number of days to skin ulceration, consistent with previous UHDR studies investigating skin response (Singers Sorensen *et al*., 2022; Duval *et al*., 2023; Pogue *et al*., 2024). The skin toxicity scoring scale was similar to that previously published by Noda et al. (Noda *et al*., 2005), shown in Table 2. As stated previously, the event conditional for a Kaplan-Meier curve analysis for this study was skin ulceration across the irradiation area, which corresponds to a skin score of 2 on this guide. This score was chosen because the observation of distinct ulceration onset was determined to be the most objectively binary decision point between graders for onset of radiation induced normal tissue toxicity. Additionally, each mouse was survived and monitored daily until a skin score of 3 was reached, at which point the mouse was sacrificed based upon IACUC guidance. At the conclusion of the study, 51 days post-irradiation, all remaining mice were sacrificed regardless of skin score. 51 days was chosen to allow for the observation of recovery in low toxicity presenting mice (Ilina *et al*., 2025). Daily reading scores for each mouse were determined with knowledge and consideration of previous daily scores to track induced toxicity progression over time, while the mouse groups (UHDR, CDR, oxygen conditions) were blinded from the grader.

**Table 2:** Radiation-induced skin toxicity numerical score scale used to quantify damage from daily photographs.

| **Skin Radiation Toxicity Score Scale** |  |
| --- | --- |
| Score | Description |
| 3 | Severe skin breakdown affecting majority of irradiated area with extensive moist exudate |
| 2.5 | Large confluent areas of moist desquamation with exudate |
| 2 | Extensive area breakdown with moist regions (ulceration) |
| 1.75 | Multiple small areas of moist desquamation |
| 1.5 | Localized moist desquamation in a very small area with scaly or crusty appearance |
| 1.25 | Pronounced erythema with white flaky skin and/or edema |
| 1 | Clear abnormality with erythema (reddening) |
| 0.75 | Mild abnormality; subtle changes are visible |
| 0.5 | Questionable change; difficult to distinguish from normal |
| 0 | Normal appearance; no visible changes |

### Tissue Oxygenation Modulation

As detailed in Figure 1A, five oxygenation conditions were used to modulate tissue oxygen levels:

i. inhaled room air with isoflurane anesthesia, with complete vascular compression to the leg above the irradiation site stopping all blood flow, (“Anoxic”)
ii. inhaled room air with isoflurane anesthesia, with partial vascular compression above the irradiation site inducing restricted blood flow, (“Hypoxic to mildly hypoxic)”
iii. room air with isoflurane anesthesia, with no leg ligation (21% O_2_), (“Mildly hypoxic”)
iv. inhaled 100% oxygen gas with isoflurane anesthesia, with no leg ligation, (“Normoxic to Mildly hyperoxic)
v. inhaled carbogen gas, here defined as 95% O_2_ with 5% CO_2_, with anesthesia, with no leg ligation. (“Hyperoxic”)

**Figure 1:**
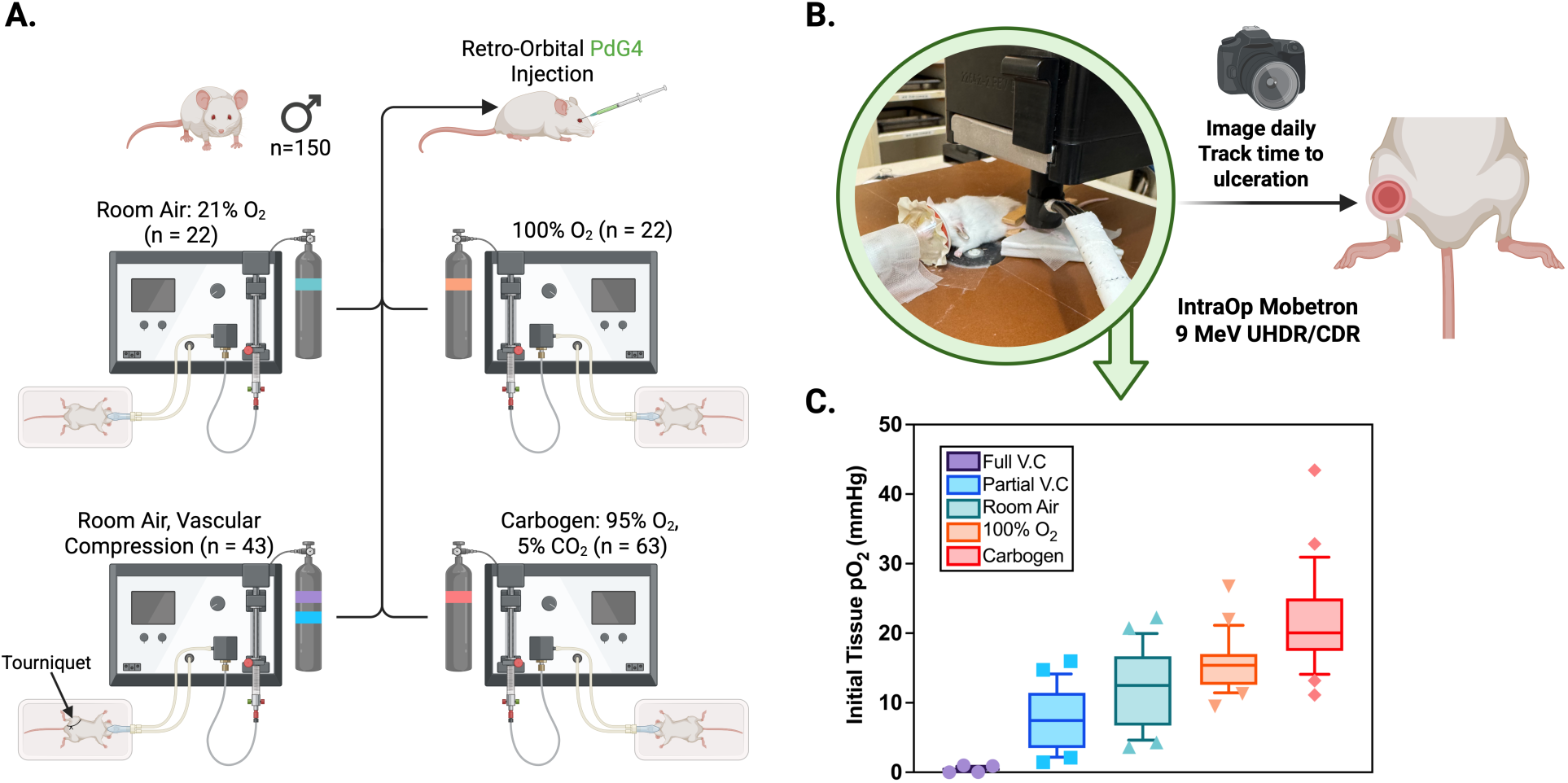
A) Study design, including population distribution, probe injection method, and oxygenation cohorts (room air, room air with vascular compression, 100% oxygen, and carbogen). Gas canisters are color-coded to match the legend and data in (C). B) Radiation delivery setup, showing mouse positioning photograph, and endpoint tracking. C) Box plots of initial tissue pO_2_ at irradiation for each oxygen condition. Full vascular compression (V.C) with room air (purple), partial vascular compression with room air (blue), inhaled room air (green), inhaled 100% oxygen (orange), and inhaled carbogen (red).

These approaches achieved a range of initial pO_2_ levels from 0 to 43 mmHg in the irradiation region of tissue on the mouse leg (Figure 1C). Both the UHDR (n = 54) and CDR (n = 55) groups were divided (n = 10-11) into these five cohorts to evaluate the FLASH effect under varying oxygen conditions, given the expected impact of oxygen levels on radiation response (Hall and Amato, 2018). An additional cohort of UHDR (n = 20) and CDR (n = 21) mice were irradiated at 20 or 15 Gy dose levels under carbogen conditions due to assay saturation.

### *In Vivo* Oximetry

Real-time tissue oxygenation monitoring was accomplished using the phosphorescence-based oxygen probe Oxyphor PdG4 (Oxygen Enterprises, Philadelphia, PA). This dendritic probe operates via oxygen-dependent phosphorescence quenching and has undergone extensive validation for both *in vitro* and *in vivo* applications (Esipova *et al*., 2011). The Oxyphor PdG4 was administered through retro-orbital injection at 0.05 mL from 200 µM stock concentration, dissolved in phosphate buffer solution, approximately 60 minutes before radiation exposure to ensure adequate vascular probe dispersion throughout the animal.

Oxygen measurements were captured using an OxyLED phosphorimeter system (Oxygen Enterprises, Philadelphia, PA). The system operated at a temporal resolution of 10 Hz (100 ms intervals), providing high-resolution oxygen dynamics both immediately prior to and following irradiation. To maintain measurement integrity while preventing radiation field interference, optical fibers were positioned at a 15 mm offset from the radiation field’s center, oriented 45 degrees to the normal so as to allow adequate measuring of the irradiated tissue without impeding the electron beam, with the main instrumentation and data acquisition computer situated approximately 1 meter from the radiation source. Measurements of phosphorescence from the skin tend to be dominated from signals within the skin, with lesser contributions to the underlying tissue, therefore we interpreted the signal as coming predominantly from the skin. Conversion of phosphorescence lifetime measurements to absolute pO_2_ values was performed using pre-measured calibration constants for this batch of Oxyphor (Esipova *et al*., 2011), calibrated at a temperature of 37°C.

This measurement approach enabled quantification of radiation-induced oxygen dynamics in real time. Baseline tissue oxygenation was established by averaging pre-irradiation measurements per mouse, which were then subtracted from values obtained during and after radiation exposure (Sunnerberg *et al*., 2025). The resulting differential oxygen measurements (ΔpO_2_) were normalized to radiation dose, yielding oxygen consumption g-values (g_O2_), expressed as mmHg per unit dose delivered (mmHg/Gy).

### Statistical Analysis

For the radiation-induced skin damage scoring, numerical values were evaluated daily with the standardized scoring system (Table 2) by trained observers. To address the issue of mice being removed from the study upon reaching endpoints, a last-observation-carried-forward (LOCF) approach was used, where the final observed skin score for each mouse was carried forward for all subsequent time points. This method preserved the impact of severe reactions in statistical analysis and prevented skewing of data as mice were removed from study. Using this dataset, a mixed-effects analysis was used to compare UHDR and CDR cohorts within each oxygen environment over the post-irradiation observation period. The tests included dose rate and time as fixed effects, with individual mice as random effects to account for correlation among repeated measures within each mouse. This approach appropriately handled the ordinal nature of the skin score data while accommodating the longitudinal study design. Separate tests were run for each oxygenation condition to determine whether dose rate significantly influenced skin toxicity progression under specific oxygen environments. The null hypothesis tested was that dose rate did not affect the progression of radiation-induced skin toxicity under equivalent oxygenation conditions. Statistical significance was defined as p-value ≤ 0.05.

To evaluate the difference in skin ulceration progression (score = 2 in Table 2), Kaplan-Meier plot analysis were used, displaying the proportion of mice that remained ulceration-free over the observation period. Separate curves were generated for UHDR and CDR treatment groups within each oxygenation condition. Statistical comparisons between dose rate groups were performed using Log-Rank (Mantel-Cox) tests, with p-value ≤ 0.05 being considered statistically significant. This analysis allowed quantitative assessment of whether dose rate affects the timing and incidence of ulceration under controlled oxygen environments.

Comparison of oxygen consumption g-values and initial pO_2_ values between mice that developed ulceration and those that remained ulceration-free was performed using Mann-Whitney U tests. The two experimental hypotheses (H_1_) tested were that pO_2_ and g_O2_ are lower for UHDR-irradiated mice that did not develop ulcers through the duration of the study. Statistical significance was defined as p-value ≤ 0.05.

## Results

### Tissue pO_2_ Observations in Treatment Groups

Tissue pO_2_ values were varied through systematic induction of breathing gas or through clamping of blood flow, as illustrated in Figure 1. The induced ranges of irradiated tissue pO_2_, with mean and standard deviations illustrated in the graph in Figure 1C, were: (i) full vascular compression breathing room air, pO_2_ = 0.44 ± 0.26 mmHg, (ii) partial vascular compression breathing room air, pO_2_ = 7.4 ± 4.3 mmHg, (iii) freely breathing room air, pO_2_ = 11.9 ± 5.6 mmHg, (iv) freely breathing 100% oxygen, pO_2_ = 15.7 ± 3.8 mmHg, and (v) freely breathing carbogen, pO_2_ = 21.4 ± 6.9 mmHg. The total ranges were: full vascular compression 0-1.00 mmHg, partial vascular compression 1.4-16.0 mmHg, room air 3.6-22.3 mmHg, 100% oxygen 9.5-26.7 mmHg, and carbogen 11.1-43.4 mmHg. These breathing and clamping groups were taken as treatment cohorts for the remaining skin toxicity response studies in order to systematically study the variation in radiation induced skin toxicity.

### Dependence of Skin Radiation Toxicity Scores on Baseline pO_2_

Radiation-induced skin toxicity was monitored daily for 51 days post-irradiation, and mean skin scores over time for each cohort are shown in Figure 2 for each of the five oxygen conditions (panels A-E), with both CDR and UHDR groups on each of these. To analyze these data, a mixed-effects analysis with LOCF was used to reveal any significant differences between UHDR and CDR treatment groups under specific oxygenation conditions. Mice in the room air cohort (Figure 2C) exhibited significantly reduced skin toxicity following UHDR treatment compared to CDR (p-value = 0.001). A similar FLASH sparing effect was observed in the room air with partial vascular compression (Figure 2B) cohort (p-value = 0.005), and to a lesser extent in the 100% oxygen (Figure 2D) cohort (p-value = 0.03), where UHDR-treated mice had less severe reactions throughout the study.

**Figure 2:**
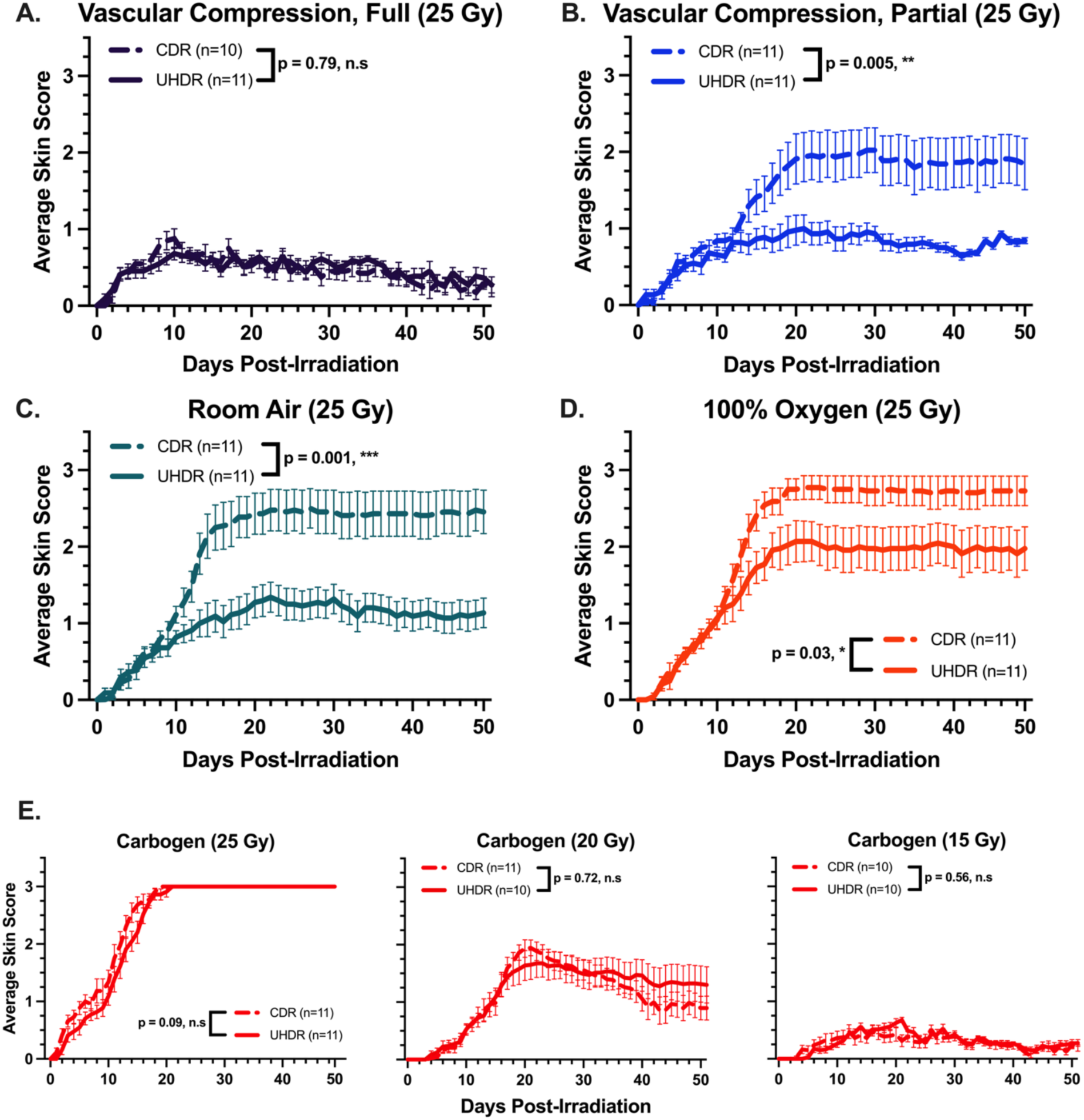
Average skin lesion progression daily for each oxygenation cohort. Solid lines represent UHDR data, and dashed lines represent CDR data. Error bars show standard error of the mean. Statistical significance of dose rate effect is indicated as ***(p-value ≤ 0.001), **(p-value ≤ 0.01), *(p-value ≤ 0.05), and n.s. (p-value > 0.05). A) Room air with complete vascular compression (purple, p-value = 0.79). B) Room air with partial vascular compression (blue, p-value = 0.005). C) Room air (green, p-value = 0.001). D) 100% oxygen (orange, p-value = 0.03). E) Carbogen inhalation for each of 25 (left), 20 (middle) and 15 (right) Gy dose cohorts (red, p-values = 0.09, 0.72, 0.56).

In contrast, no significant differences were observed between UHDR and CDR groups in the carbogen breathing cohorts. Mice in the 25 Gy carbogen (p-value = 0.09) group developed comparable skin toxicities, with assay saturation reached at 3.0. Additional mouse cohorts were treated with carbogen at 20 and 15 Gy dose levels (Figure 2 E, middle and right), and in neither of these was a significant FLASH skin sparing effect observed (p-values = 0.72 & 0.56), despite not reaching assay score saturation.

Kaplan-Meier curves were plotted for the mice, as shown in Figure 3, illustrating the differences in ulceration between UHDR and CDR groups across oxygenation conditions. In the normoxic (room air) cohort, UHDR reduced overall progression to ulceration compared to CDR (p-value = 0.01, Figure 3C), with 8 out of 11 UHDR-treated mice remaining ulcer-free for the full study duration. In contrast, the median time to ulceration for CDR-treated mice was 15 days.

**Figure 3:**
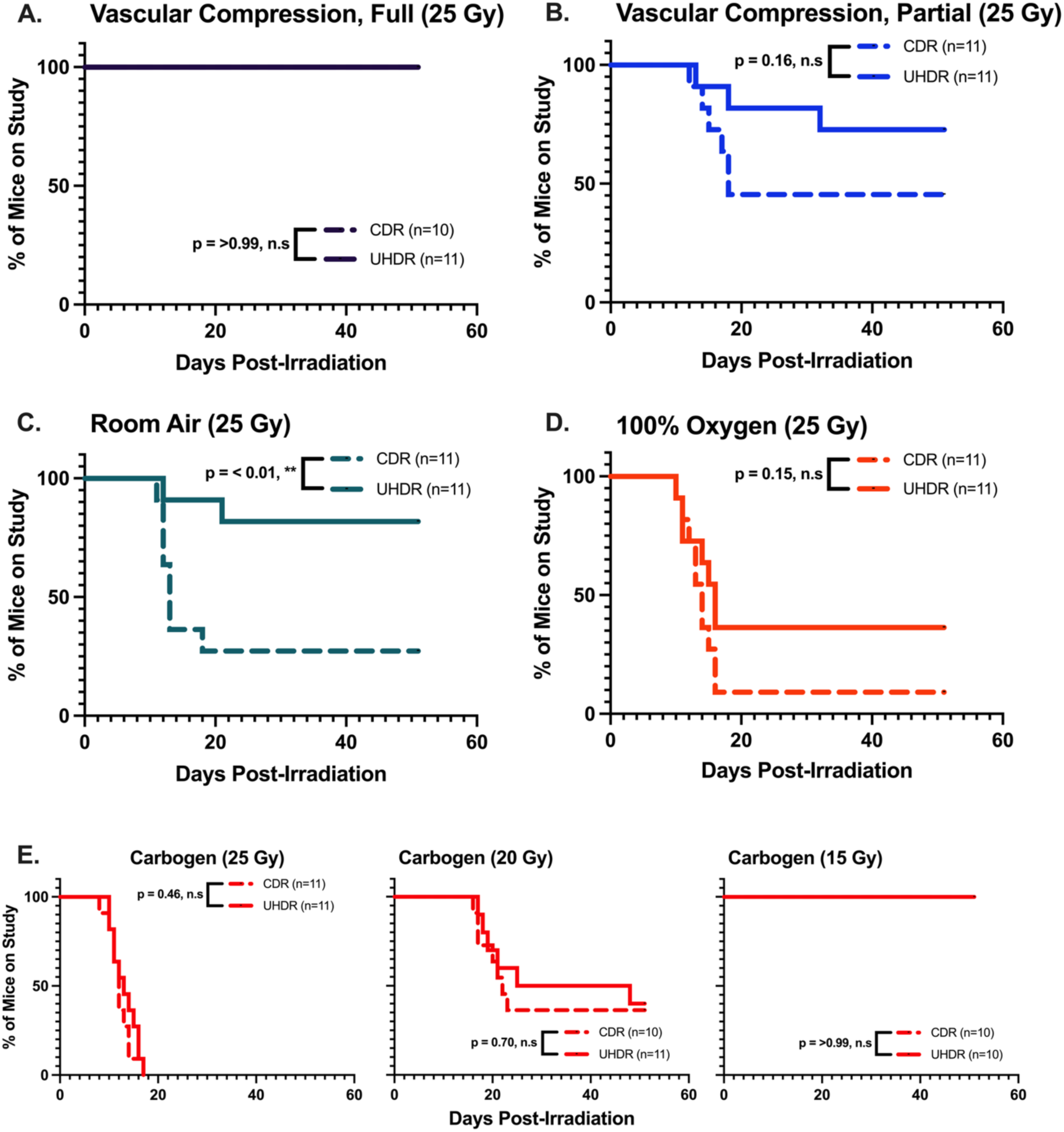
Kaplan-Meier curves for ulcer-free progression of mice in each oxygenation cohort. Solid lines represent UHDR data, and dashed lines represent CDR data. Statistical significance of dose rate effect is indicated as ** (p-value ≤ 0.01), * (p-value ≤ 0.05), and n.s. (p-value > 0.05). A) room air with full vascular compression (purple, p-value = >0.99). B) Room air with partial vascular compression (blue, p-value = 0.16). C) Room air (green, p-value = < 0.01). D) 100% oxygen (orange, p-value = 0.15). E) Carbogen groups given 25 (left), 20 (middle) and 15 (right) Gy, respectively (red, p-value = 0.46, 0.70, >0.99).

No significant difference was seen in ulceration-free progression between UHDR and CDR groups in the 100% oxygen (Figure 3D, p-value = 0.15) cohort. Similarly, no significant difference was observed with any of the carbogen groups at 25, 20 or 15 Gy (Figure 3E). This range of doses was specifically used to test the full response range of damage versus dose when Carbogen was used.

Despite a significant difference in skin scores (Figure 2B, p-value = 0.005), the partial vascular compression cohort did not show a statistically significant FLASH effect for ulceration-free progression (Figure 3B, p-value = 0.16). This likely reflects an overall reduction in radiation-induced toxicity for CDR-treated mice in this cohort, leading to fewer ulceration events and in turn an underpowered log-rank test.

To illustrate the observed skin radiation toxicity dependency upon baseline tissue pO_2_, the average skin toxicity scores at the point of maximum damage (20 days post irradiation) were plotted for all five cohorts (full vascular compression, partial vascular compression, room air, 100% O_2_, and carbogen) against the measured mean initial tissue pO_2_, in Figure 4. A non-linear fit of a sigmoid function, a four-parameter logistic curve, was additionally fit to the data to illustrate the trend for separation in toxicity response between UHDR and CDR cohorts. Maximum separation between UHDR and CDR toxicity occurred under initial tissue oxygen tensions between 7 and 15 mmHg.

**Figure 4:**
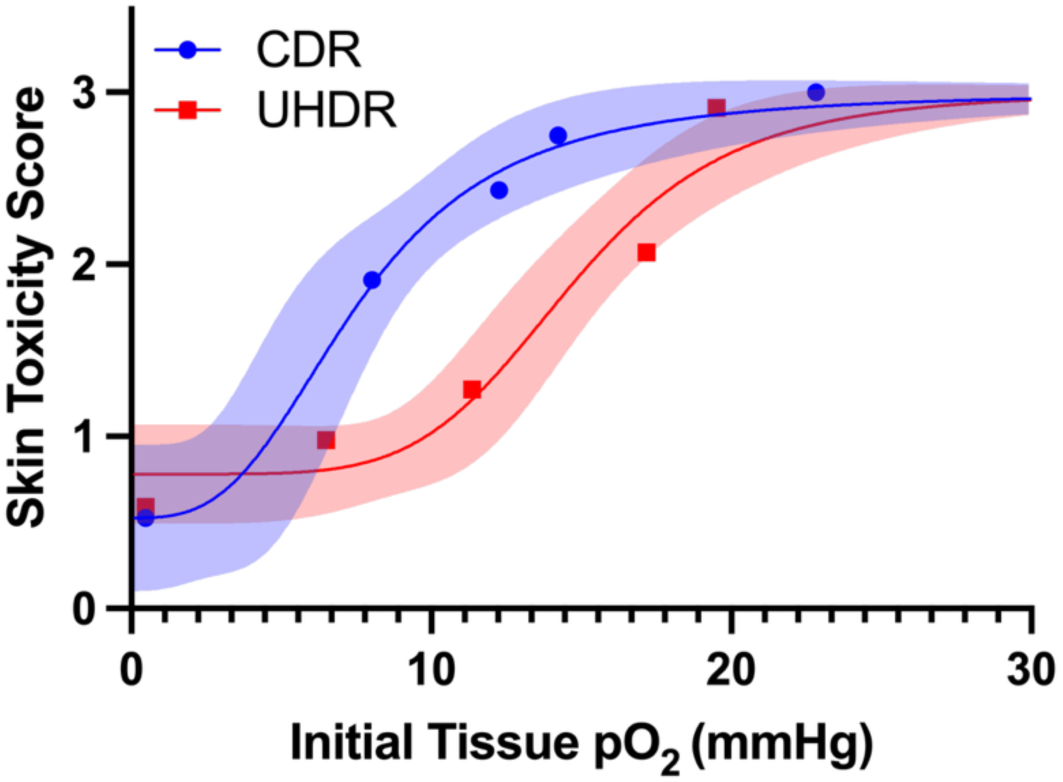
Mean skin toxicity scores on day 20 versus mean initial tissue pO_2_ are plotted for the 5 treatment groups at 25 Gy dose, for CDR (red) and UHDR (blue). Non-linear sigmoid curves are fit to the data, with 95% confidence bands in shaded color.

### Correlating Baseline pO_2_ and g_O2_ With Radiation Induced Ulceration

The relationship between initial tissue oxygen tension (pO_2_), oxygen consumption (g_O2_), and ulceration is shown in Figure 5, with the raw data plotted in Figure 5A, separating those animals that showed ulceration (red points) from those that did not (blue points). Three samples of oxygen tension measurements during irradiation are shown in Figure 5D, with depletion events highlighted in red. Among UHDR-irradiated mice, those that developed ulceration had significantly higher g_O2_-values (0.17 ± 0.06 vs. 0.03 ± 0.07 mmHg/Gy, p << 0.001) and initial pO_2_ levels (16.9 ± 5 vs. 3.8 ± 5.4 mmHg, p-value << 0.001) as compared to mice that remained skin ulcer-free, as shown in Figures 5B and 5C. An exponential functional regression fit of g_O2_ and pO_2_ yielded an R^2^ value of 0.86.

**Figure 5:**
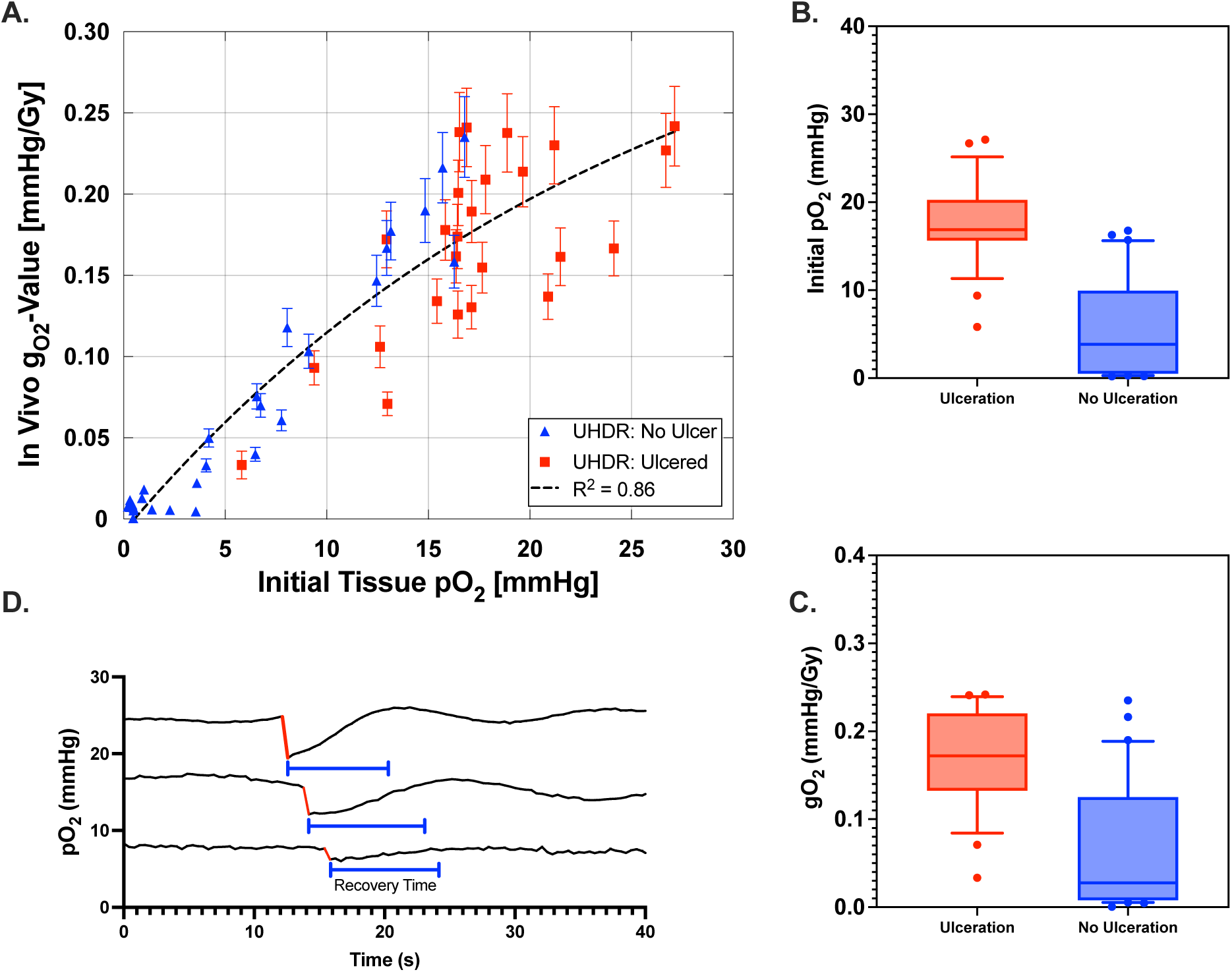
**A)** In vivo g_O2_ measurements for UHDR-irradiated mice plotted against initial tissue pO_2_. Each data point represents an individual mouse, with ulcered mice shown as red squares and non-ulcered mice as blue triangles. B) Box and whisker plots for initial pO_2_ measurements between ulcerating (red) and non-ulcerating (blue) mice. C) Box and whisker plots for gO_2_ measurements between ulcerating (red) and non-ulcerating (blue) mice. D) Sample recordings of murine tissue pO_2_ using the OxyLED system during UHDR irradiation, showing oxygen consumption (red) within the irradiated field.

## Discussion

The FLASH effect of reducing normal tissue toxicity under ultra-high dose rates is well established, though the role that oxygen plays in modulating this effect, along with its mechanism, remains demonstrated in only a few studies and at isolated oxygen points. *In vitro* data has been more systematic because it is easier to control oxygen, and these studies have shown a FLASH sparing effect present at low-range oxygen values (2-4 mmHg), disappearing under higher oxygen values (24 mmHg) (Cooper *et al*., 2022). *In vivo* data has shown that UHDR irradiations achieve equivalent, and possibly superior, control in clamped tumors held at anoxia, when compared to CDR cohorts (Leavitt *et al*., 2024). A recent comprehensive study of NTCP damage response across the full range of doses was completed in skin FLASH sparing, quantifying how tissue clamping to stop blood flow, inducing anoxia, reduces the dose modifying factor from 1.43 in normal to 0.96 in clamped tissue (Hansen *et al*., 2025). Another recent study showed that the NTCP data showed no differences in Dose Modifying Factor (DMF) when mice were breathing air (DMF=1.42) or oxygen (DMF=1.40) (Sesink *et al*., 2026). Interestingly, though, this paper also showed in colon that there was a loss of the FLASH sparing effect under oxygen breathing (100% O_2_) versus air (21% O_2_). The first study to explore FLASH at hyperoxia in the rat brain model showed a loss of the FLASH protective effect when supplemental oxygen was mixed with anesthesia gas (isoflurane) at the time of irradiation (Iturri *et al*., 2023). Thus the trends of tissue radiation induced toxicity response with changes in oxygen and blood flow are a complex landscape.

The present study was designed to test skin radiation toxicity *in vivo* via systematic variation in tissue oxygen, varied across the fullest range from complete anoxia up to hyperoxia with carbogen, while comparing CDR and UHDR groups at a fixed moderate dose. The study results demonstrate that observable differences in radiation induced toxicity scores exist between CDR and UHDR in this *in vivo* skin model at the dose level evaluated, and we term this difference as a single dose estimate of FLASH skin sparing. The sparing is clearly dependent on initial tissue oxygen tension at the 25 Gy dose level investigated, being present in the ranges of oxygen found in the three conditions of room air, 100% oxygen and partial vascular compression (pO_2_ ≈ 4-18 mmHg). However, when oxygen levels were modulated – either by complete anoxia (complete leg compression with ligation) inducing hypoxia (pO_2_ ≈ 0 mmHg) or hyperoxia, via carbogen inhalation (pO_2_ ≈ 15-28 mmHg), significant differences between UHDR and CDR induced toxicity were not observed. This is illustrated in Figure 4, where UHDR irradiation is seen to have suppressed damage at mid-oxygen levels, as compared to CDR cohorts, in the range of pO_2_ ≈ 7-16 mmHg. It is worth noting that the shape of the sigmoidal line fit for CDR data matches the classically illustrated oxygen enhancement ratio (OER) curve (Grimes and Partridge, 2015; Hall and Amato, 2018). With this analogy, UHDR appears to reduce the effect of oxygen mediated damage in this intermediate oxygen range, or it could be that the damage is shifted right to occur at higher oxygen levels. An alternate interpretation is that the mechanism of FLASH sparing results from a reduction in oxygen-mediated damage that is not accessible at moderate pO_2_ values. The cause of a loss of FLASH sparing at high and low pO_2_ values could potentially be from different mechanisms and could even be more related to changes in blood flow than oxygen. Further research into these hypotheses is necessary. Taken together, these finding suggest that a baseline oxygenation level is likely a critical factor influencing not just the observation of the UHDR FLASH sparing effect but is also likely, at least, part of the mechanism causing the effect.

Examining the results from the hyperoxia carbogen group, it was noted that the *in vivo* assay saturated at the highest toxicity level for 25 Gy, with all mice reaching the maximum skin score of 3.0 in both UHDR and CDR groups (Figure 2E). This skin toxicity scoring scale saturates at a maximum of 3.0 and so a range of dose groups at 20 and 15 Gy were also included in the study under carbogen to test for systematic damage shifts across the doses at an elevated tissue pO_2_ (Figure 2E). However as seen in Figures 2E and 3E, there was no significant FLASH sparing observable when these lower dose levels were used. It is possible that finer measurement techniques of toxicity reduction may yield an observation of differences under these conditions, but the magnitude of sparing does not seem to match that seen under partial vascular compression, room air and 100% oxygen breathing conditions. Thus, these data are consistent with a hypothesis that excess oxygen or increased blood flow may negate the FLASH effect, suggesting that perhaps the FLASH effect itself is dependent upon some spatial or temporal limitation in oxygen at the irradiation site, consistent with previous studies (Montay-Gruel *et al*., 2019; Adrian *et al*., 2020; Petersson *et al*., 2020; Cao *et al*., 2021; Jansen *et al*., 2021; Van Slyke *et al*., 2022; Grilj *et al*., 2024; Tavakkoli *et al*., 2024; Sunnerberg *et al*., 2026). Carbogen breathing is known to acidify tissues and lower blood and extracellular pH though the effects of this pH shift on normal tissue sparing under UHDR and CDR conditions remains to be investigated, and its impact on the FLASH effect remains unknown.

The link between oxygen tension and radiation induced tissue toxicity appears central to the mechanisms of FLASH sparing, in terms of the underlying radiation chemistry, and it has been hypothesized that the oxygen consumption itself is leading to sparing. Under UHDR conditions, the relationship between pO_2_ and tissue toxicity appears to be altered such that the expected severity of induced toxicity following 25 Gy is reduced for all but anoxic and hyperoxic tissues, showing the FLASH effect is present. In the UHDR setting, oxygen is consumed rapidly in the formation of peroxyl radicals and superoxide species, thereby causing fixation of the radiation induced damage in proteins and DNA (Grimes and Partridge, 2015; Cooper *et al*., 2022; Sunnerberg *et al*., 2023). This occurs on a short time scale, significantly shorter than under CDR conditions, where diffusion of oxygen back into the irradiated site occurs at a rate slower than consumption. It has been previously shown *in vitro* that UHDR irradiation reduces oxygen consumption as compared to CDR, which may supporting the hypothesis that UHDR leads to less oxidative modification of DNA by peroxyl radicals (Sunnerberg *et al*., 2023). However, linking *in vitro* data and DNA data to *in vivo* observations of radiation induced toxicity is inherently challenging and at best, via a modification of inhaled gas mixtures, a correlative link can be established, rather than causative, to a proposed mechanism. Still the data here is consistent with the idea that oxygen influences the presentation of FLASH normal tissue sparing, as it pertains to skin toxicity. Many studies have postulated and modeled transient oxygen depletion as a hypothesized mechanism for reduced toxicity under UHDR (Favaudon, Labarbe and Limoli, 2022), although direct *in vivo* measurements of the level of consumption, here and in previous work (Cao *et al*., 2021; El Khatib *et al*., 2022; Grilj *et al*., 2024), indicate that the level of oxygen consumed would not likely be sufficient for complete ‘depletion’ of local oxygen. Thus, there is not a clear explanation for this discrepancy and further data may be needed to determine if there are micro-localized depletions of oxygen or oxygen-related effects that are causative of reduced ROS damage *in vivo*.

Figure 2A, relative to the other panels (Figure 2B, 2C, and 2D), provides important insight into the role and potential mechanism of FLASH occurrence, based upon the loss of any sparing under anoxic conditions. In this anoxic cohort, overall skin toxicity was reduced compared to room air, and there was no difference in toxicity between UHDR and CDR groups. This observation could suggest that whatever is causing the FLASH tissue sparing must rely upon oxygen being present in the tissue. However, it is important to recognize that the dose level investigated (25 Gy) may have been too low to elicit an observable damage differential between UHDR and CDR cohorts under anoxic conditions. Further work should evaluate whether the FLASH sparing effect persists at these anoxic oxygen levels, which may be relevant for future explorations into tumors and the FLASH normal tissue sparing effect.

Overall, these results suggest that there is a need to carefully control and report tissue oxygenation conditions when designing preclinical FLASH studies, and this can be as subtle as causing variations in tissue oxygen from breathing gas, animal body temperature or mouse positioning (i.e. leg clamping). The recent data from Hansen et al has shown that the DMF effect of oxygenated versus anoxic tissue can be double the DMF of the FLASH effect itself in normoxic tissue. Thus, variation in oxygen will inevitably lead to both real and artificial changes in an observed FLASH effect if uncontrolled or unmeasured. The work here contributes to a more robust understanding of how oxygen interacts with dose rate variation in radiation skin toxicity. Future studies might extend these observations to other tissue types and dose rate regimes, further exploring how oxygen consumption and radical chemistry at UHDR influence both acute and long-term normal tissue outcomes. Further, studies that separate the independent but coupled effect of blood flow from tissue oxygenation could be very important to understand the underlying mechanisms of how clamping and carbogen affect radiation sensitivity at UHDR. Understanding how UHDR effectively influences the OER across varied tissues will be critical for refining the mechanistic basis of the FLASH effect and for optimally translating these findings into clinical practice.

## Conclusion

This study demonstrated that in murine skin, tissue oxygen tensions achieved under room air anesthetic conditions are required for the most apparent observation of a skin toxicity reduction effect from UHDR versus CDR irradiation. Under partial vascular compression (hypoxic to mildly hypoxic), room air (mildly hypoxic) and 100% oxygen (normoxic to mildly hyperoxic) conditions, UHDR treatment resulted in a significant reduction in overall radiation induced tissue toxicity compared to conventional dose rate exposures, and only the partial vascular compression condition achieved a significant reduction in overall ulceration. Hyperoxic skin with elevated local oxygen tension from carbogen breathing animals achieved maximal toxicity, and did not demonstrate any UHDR induced FLASH sparing, at any dose level studied. Anoxic skin from leg clamping inducing complete blood flow occlusion exhibited lower levels of radiation induced toxicity and no dose rate dependent sparing effect. Measures of baseline pO_2_ and radiolytic consumption, g_O2_, during UHDR irradiation confirmed that they were related, especially in non-ulcered skin at lower pO_2_, and both were significantly related to the likelihood of skin ulceration at the pO_2_ levels studied here.

These findings suggest that the oxygen-mediated enhancement of radiation damage that is conventional in radiobiology is suppressed when UHDR is used, as seen in mouse skin. Oxygen appears therefore to be a primary modulator of FLASH, because this sparing effect is only present at low-to-middle levels of oxygen tension in skin. Anoxia from tissue clamping appears to negate dose rate differences in skin toxicity, perhaps indicating that the absence of oxygen leads to a dose rate independent mechanism, as has been documented in tumors (Leavitt *et al*., 2024). Additionally, carbogen increases blood flow and induces hyperoxic effects that may negate toxicity differences between UHDR and CDR, suggesting that there is an ideal oxygen value for maximal FLASH sparing of a given tissue.

## Acknowledgements

Oxyphor PdG4 was supplied through collaboration with Professor Sergei Vinogradov at the University of Pennsylvania.

## Data Availability Statement

Research data will be shared upon reasonable request to the corresponding author.

## References

Adrian, G. et al. (2020) “The FLASH effect depends on oxygen concentration,” The British Journal of Radiology, 93(1106), p. 20190702. Available at: 10.1259/bjr.20190702.

Cao, X. et al. (2021) “Quantification of Oxygen Depletion During FLASH Irradiation In Vitro and In Vivo,” International Journal of Radiation Oncology Biology Physics. Available at: 10.1016/j.ijrobp.2021.03.056.

Colizzi, I. et al. (2026) “Systematic analysis of biological endpoint variability and implications for quantitative modeling of the FLASH sparing effect,” Physics and Imaging in Radiation Oncology, 37, p. 100915. Available at: 10.1016/j.phro.2026.100915.

Cooper, C.R. et al. (2022) “FLASH irradiation induces lower levels of DNA damage ex vivo, an effect modulated by oxygen tension, dose, and dose rate,” Br J Radiol. Available at: 10.1259/bjr.20211150.

Dewey, D.L. and Boag, J.W. (1959) “Modification of the Oxygen Effect when Bacteria are given Large Pulses of Radiation,” Nature, 183(4673), pp. 1450–1451. Available at: 10.1038/1831450a0.

Duval, K.E.A. et al. (2023) “Comparison of Tumor Control and Skin Damage in a Mouse Model after Ultra-High Dose Rate Irradiation and Conventional Irradiation,” Radiation Research. Available at: 10.1667/rade-23-00057.

El Khatib, M., et al. (2022) “Ultrafast Tracking of Oxygen Dynamics During Proton FLASH,” Int J Radiat Oncol Biol Phys. Available at: 10.1016/j.ijrobp.2022.03.016.

El Khatib, M., et al. (2024) “Direct Measurements of FLASH-Induced Changes in Intracellular Oxygenation,” International Journal of Radiation Oncology, Biology, Physics, 118(3), pp. 781–789. Available at: 10.1016/j.ijrobp.2023.09.019.

Esipova, T.V. et al. (2011) “Two New ‘Protected’ Oxyphors for Biological Oximetry: Properties and Application in Tumor Imaging,” Analytical chemistry, 83(22), pp. 8756–8765. Available at: 10.1021/ac2022234.

Favaudon, V. et al. (2014) “Ultrahigh dose-rate FLASH irradiation increases the differential response between normal and tumor tissue in mice,” Science Translational Medicine, 6(245), p. 245ra93. Available at: 10.1126/scitranslmed.3008973.

Favaudon, V., Labarbe, R. and Limoli, C.L. (2022) “Model studies of the role of oxygen in the FLASH effect,” Med Phys. Available at: 10.1002/mp.15129.

Grilj, V. et al. (2024) “In vivo measurements of change in tissue oxygen level during irradiation reveal novel dose rate dependence,” Radiother Oncol. Available at: 10.1016/j.radonc.2024.110539.

Grimes, D.R. and Partridge, M. (2015) “A mechanistic investigation of the oxygen fixation hypothesis and oxygen enhancement ratio,” Biomed Phys Eng Express. Available at: 10.1088/2057-1976/1/4/045209.

Ha, B. et al. (2022) “Real-time optical oximetry during FLASH radiotherapy using a phosphorescent nanoprobe,” Radiotherapy and Oncology: Journal of the European Society for Therapeutic Radiology and Oncology, 176, pp. 239–243. Available at: 10.1016/j.radonc.2022.08.011.

Hall, E.I. and Amato, I.G. (2018) “Radiobiology for the Radiologist.” Wolters Kluwer.

Hansen, A.H. et al. (2025) “Impact of oxygen deprivation on the FLASH effect for skin toxicity in a murine model,” Radiotherapy and Oncology: Journal of the European Society for Therapeutic Radiology and Oncology, 214, p. 111277. Available at: 10.1016/j.radonc.2025.111277.

Hansen, A.H. et al. (2026) “Impact of oxygen deprivation on the FLASH effect for skin toxicity in a murine model,” Radiotherapy and Oncology: Journal of the European Society for Therapeutic Radiology and Oncology, 214, p. 111277. Available at: 10.1016/j.radonc.2025.111277.

Hornsey, S. and Bewley, D.K. (1971) “Hypoxia in Mouse Intestine Induced by Electron Irradiation at High Dose-rates,” International Journal of Radiation Biology and Related Studies in Physics, Chemistry and Medicine, 19(5), pp. 479–483. Available at: 10.1080/09553007114550611.

Ilina, A. et al. (2025) “FLASH effect is diminished by daily fractionation of electron RT in mouse skin,” Physics in Medicine and Biology, 70(23), p. 235020. Available at: 10.1088/1361-6560/ae205e.

Iturri, L. et al. (2023) “Oxygen supplementation in anesthesia can block FLASH effect and anti-tumor immunity in conventional proton therapy,” Commun Med (Lond). Available at: 10.1038/s43856-023-00411-9.

Jansen, J. et al. (2021) “Does FLASH deplete oxygen? Experimental evaluation for photons, protons, and carbon ions,” Medical Physics, 48(7), pp. 3982–3990. Available at: 10.1002/mp.14917.

Jansen, J. et al. (2022) “Changes in Radical Levels as a Cause for the FLASH effect: Impact of beam structure parameters at ultra-high dose rates on oxygen depletion in water,” Radiotherapy and Oncology: Journal of the European Society for Therapeutic Radiology and Oncology, 175, pp. 193–196. Available at: 10.1016/j.radonc.2022.08.024.

Kirby-Smith, J.S. and Dolphin, G.W. (1958) “Chromosome Breakage at High Radiation Dose-Rates,” Nature, 182(4630), pp. 270–271. Available at: 10.1038/182270a0.

Leavitt, R.J. et al. (2024) “Acute Hypoxia Does Not Alter Tumor Sensitivity to FLASH Radiation Therapy,” Int J Radiat Oncol Biol Phys. Available at: 10.1016/j.ijrobp.2024.02.015.

Montay-Gruel, P. et al. (2017) “Irradiation in a flash: Unique sparing of memory in mice after whole brain irradiation with dose rates above 100Gy/s,” Radiotherapy and Oncology: Journal of the European Society for Therapeutic Radiology and Oncology, 124(3), pp. 365–369. Available at: 10.1016/j.radonc.2017.05.003.

Montay-Gruel, P. et al. (2019) “Long-term neurocognitive benefits of FLASH radiotherapy driven by reduced reactive oxygen species,” Proceedings of the National Academy of Sciences of the United States of America, 116(22), pp. 10943–10951. Available at: 10.1073/pnas.1901777116.

National Research Council (US) Committee on the Biological Effects of Ionizing Radiation (BEIR V) (1990) Health Effects of Exposure to Low Levels of Ionizing Radiation: Beir V. Washington (DC): National Academies Press (US). Available at: http://www.ncbi.nlm.nih.gov/books/NBK218704/ (Accessed: December 4, 2025).

Noda, S. et al. (2005) “Inter-strain variance in late phase of erythematous reaction or leg contracture after local irradiation among three strains of mice,” Cancer Detection and Prevention, 29(4), pp. 376–382. Available at: 10.1016/j.cdp.2005.06.005.

Petersson, K. et al. (2020) “A Quantitative Analysis of the Role of Oxygen Tension in FLASH Radiation Therapy,” International Journal of Radiation Oncology, Biology, Physics, 107(3), pp. 539–547. Available at: 10.1016/j.ijrobp.2020.02.634.

Pogue, B.W. et al. (2024) “Major contributors to FLASH sparing efficacy emerge from murine skin studies: dose rate, total dose per fraction, anesthesia and oxygenation,” Front Oncol. Available at: 10.3389/fonc.2024.1414584.

Rahman, M. et al. (2023) “Characterization of a diode dosimeter for UHDR FLASH radiotherapy,” Medical Physics, 50(9), pp. 5875–5883. Available at: 10.1002/mp.16474.

Sesink, A. et al. (2026) “Impact of increased oxygen concentration on the FLASH sparing effect in mice is tissue dependent,” British Journal of Radiology, 99(1178), pp. 254–262. Available at: 10.1093/bjr/tqaf290.

Singers Sorensen, B., et al. (2022) “In vivo validation and tissue sparing factor for acute damage of pencil beam scanning proton FLASH,” Radiother Oncol. Available at: 10.1016/j.radonc.2021.12.022.

Sunnerberg, J.P. et al. (2023) “Mean dose rate in ultra-high dose rate electron irradiation is a significant predictor for O2consumption and H2O2yield,” Physics in Medicine and Biology, 68(16), p. 165014. Available at: 10.1088/1361-6560/ace877.

Sunnerberg, J.P. et al. (2025) “Oxygen Consumption In Vivo by Ultra-High Dose Rate Electron Irradiation Depends Upon Baseline Tissue Oxygenation,” International Journal of Radiation Oncology, Biology, Physics, 121(4), pp. 1053–1062. Available at: 10.1016/j.ijrobp.2024.10.018.

Sunnerberg, J.P. et al. (2026) “Timescale of FLASH Sparing Effect Determined by Varying Temporal Split of Dose Delivery in Mice,” International Journal of Radiation Oncology*Biology*Physics, 124(3), pp. 831–841. Available at: 10.1016/j.ijrobp.2025.09.052.

Tavakkoli, A.D. et al. (2024) “Anesthetic Oxygen Use and Sex Are Critical Factors in the FLASH Sparing Effect,” Advances in Radiation Oncology, 9(6), p. 101492. Available at: 10.1016/j.adro.2024.101492.

Tessonnier, T. et al. (2024) “Diamond detectors for dose and instantaneous dose-rate measurements for ultra-high dose-rate scanned helium ion beams,” Medical Physics, 51(2), pp. 1450–1459. Available at: 10.1002/mp.16757.

Van Slyke, A.L. et al. (2022) “Oxygen Monitoring in Model Solutions and In Vivo in Mice During Proton Irradiation at Conventional and FLASH Dose Rates,” Radiation Research, 198(2), pp. 181–189. Available at: 10.1667/RADE-21-00232.1.

Vaupel, P., Flood, A.B. and Swartz, H.M. (2021) “Oxygenation Status of Malignant Tumors vs. Normal Tissues: Critical Evaluation and Updated Data Source Based on Direct Measurements with pO2 Microsensors,” Applied Magnetic Resonance, 52(10), pp. 1451–1479. Available at: 10.1007/s00723-021-01383-6.

Wardman, P. (2007) “Chemical radiosensitizers for use in radiotherapy,” Clin Oncol (R Coll Radiol). Available at: 10.1016/j.clon.2007.03.010.

